# Comparative infection and pathogenesis of SARS-CoV-2 Omicron and Delta variants in aged and young Syrian hamsters

**DOI:** 10.1101/2022.03.02.482662

**Authors:** Nadia Storm, Nicholas A. Crossland, Lindsay G. A. McKay, Anthony Griffiths

## Abstract

Coronavirus disease 2019 continues to batter the world with the unceasing introduction of new variants of the causative virus, SARS-CoV-2. In order to understand differences in disease caused by variants of concern and to develop variant-specific vaccines, suitable small animal models are required that mimic disease progression in humans at various stages of life. In this study, we compared the dynamics of infection with two SARS-CoV-2 variants of concern (Delta and Omicron) in aged (>1 year 3 months old) and young (<5 weeks old) Syrian hamsters (*Mesocricetus auratus*). We show that no weight loss occurred in Omicron infected groups regardless of age, while infection with the Delta variant caused weight loss of up to 10% by day 7 post-infection with slower and incomplete recovery in the aged group. Omicron replicated to similar levels as Delta in the lungs, trachea and nasal turbinates, with no significant differences in the tissue viral loads of aged versus young animals for either variant. In contrast to rare necrosis observed in Omicron-infected animals regardless of age, severe necrosis was observed in the olfactory epithelium in Delta-infected animals. Omicron infection also resulted in mild pulmonary disease in both young and aged animals compared to the moderate acute necrotizing bronchointerstitial pneumonia seen in Delta-infected animals. These results suggest that Omicron infection results in an attenuated clinical disease outlook in Syrian hamsters compared to infection with the Delta variant irrespective of age.

## Introduction

Coronavirus disease 2019 (COVID-19) is a respiratory illness caused by the severe acute respiratory syndrome coronavirus 2 (SARS-CoV-2) that emerged for the first time at the end of 2019 to cause a devastating and unrelenting pandemic (Zhu *et al*., 2020). By 2021, several variants of the virus had been discovered and contributed to repeated cycles of infection in different parts of the world. In late November 2021, South Africa reported the detection of a new variant of SARS-CoV-2, B.1.1.529, characterized by more than 50 mutations compared to the reference strain, with at least 30 occurring in the spike (S) protein alone (Martin *et al*., 2022; World Health Organization, 2021). The World Health Organization soon after named the variant Omicron and classified it as a variant of concern due to its potential detrimental effect on the ongoing COVID-19 pandemic (World Health Organization, 2021(a)). The Omicron variant is associated with a reduction in neutralization by antibody treatments and post-vaccination sera, with immune escape occurring even amongst those previously infected and/or vaccinated against the virus (Cao *et al*., 2021; Dejnirattisai *et al*., 2022; Kuhlman *et al*., 2022; Muik *et al*., 2022). Since its discovery, Omicron has spread exponentially and has outcompeted Delta (B.1.617.2), another SARS-CoV-2 variant of concern that was dominating infections worldwide at the time of Omicron’s emergence. The SARS-CoV-2 Delta variant was first discovered in India late in 2020 and is characterized by at least 20 mutations of which eight occur in the spike protein (Latif *et al*., 2022). The Delta variant is associated with high transmission rates as well as increased morbidity and mortality compared to earlier SARS-CoV-2 variants (World Health Organization, 2021(b)).

Infection with SARS-CoV-2 results in a variety of clinical signs and symptoms, ranging from anosmia, fever, cough and fatigue to more severe respiratory symptoms including life-threatening pneumonia and multisystem inflammatory syndrome (Guan *et al*., 2020). Clinical disease caused by the Omicron variant appears to be less severe than disease caused by the Delta variant in humans (Iuliano *et al*., 2022). The severity of disease caused by both variants also correlates with differences in age and the presence or absence of co-morbidities, with younger, healthier people more likely to be asymptomatically infected or report mild symptoms, and obesity, hypertension, diabetes, cardiovascular disease, and old age correlating with increased risk of mortality (Zhou *et al*., 2020; Tajbakhsh *et al*., 2021; Wolter *et al*., 2022).

The Syrian (golden) hamster (*Mesocricetus auratus*) has been used widely as a model for COVID-19 due to the similar pathological features in the lungs of these animals infected with SARS-CoV-2 compared to human COVID-19 patients (Imai *et al*., 2020). Understanding the infectivity and pathogenesis of SARS-CoV-2 variants in animal models of different age ranges could assist in informing vaccine and therapeutic development strategies. To this end, we sought to determine the age-related variances in pathogenicity of infection with one of two SARS-CoV-2 variants of concern (Omicron and Delta) in the Syrian hamster model of COVID-19.

## Materials and Methods

### Ethics Clearance and Regulatory Approvals

This experiment was conducted in the animal biosafety level 4 (ABSL-4) laboratory of the National Emerging Infectious Diseases Laboratories of Boston University. Approval for this study was obtained from the Boston University Institutional Biosafety Committee (IBC) and Institutional Animal Care and Use Committee (IACUC).

### Animal husbandry and group assignment

This study consisted of 47 Syrian hamsters assigned to two main age-specific groups, with 20 aged hamsters between the ages of 15 to 16 months, and 27 young hamsters between four to five weeks of age. Each group was further divided into three sub-groups, specifically a control group, an Omicron challenge group and a Delta challenge group (for the aged group, n = 6, 7 and 7 respectively, while the young group contained nine animals per sub-group). The study animals were evaluated by a veterinarian prior to study enrollment to confirm that they were free from obvious clinical signs of ill health and were subsequently acclimated to containment for 72 hours before inoculation. The hamsters were singly housed in microisolator cages with environmental enrichment and were given food and water *ad libitum*. The study lasted for 14 days post-inoculation (DPI), with a subset of animals in each sub-group euthanized at 3 DPI (four animals in each young sub-group, and three animals in each aged sub-group except for the control group where only two were euthanized on this day).

### Challenge viruses and inoculation

Two SARS-CoV-2 variants of concern, Omicron (B.1.1.529) and Delta (B.1.617), were obtained from BEI Resources (NR-56462 and NR-55673, respectively) and were used as challenge viruses in this study. Each animal in the challenge groups was sedated with isoflurane and received 100 µl of Dulbecco’s phosphate buffered saline (DPBS; Gibco, New York, USA) containing 4.4 × 10^5^ TCID_50_/ml of either of the challenge viruses intranasally, with the dose split between the nares. Each animal in the control groups received 100 µl of DPBS only intranasally. The exposure dose was verified using a semi-solid Avicel plaque assay as described previously (Honko *et al*., 2020).

### Clinical Observations

Morbidity is not expected in the hamster model of COVID-19, therefore endpoints for this study included moribundity (respiration, appearance, responsiveness, behavior) and weight loss exceeding humane guidelines. Clinical observations were made daily to evaluate moribundity and weight loss from the day of challenge until study termination. A baseline body weight was recorded for each animal prior to inoculation, and a terminal body weight was recorded at euthanization.

### Sampling

Blood was collected from the animals at 14 DPI. Whole blood was collected from either the saphenous or gingival vein into a plastic serum separator tube and inverted 10 times. The blood samples were allowed to clot for 30 minutes at 25°C. The samples were then centrifuged at 8 000 x *g* for 3 minutes at 25°C. The serum was separated from the clot within 1 hour of collection and stored at −80°C until further processing. At euthanasia either on day 3 PI or at study termination, animals were necropsied, and the lung, trachea and nasal turbinates collected for determination of viral load and/or histopathological analysis.

### Plaque reduction neutralization test

Neutralizing antibody titers were determined in sera collected from hamsters at 14 DPI using a plaque reduction neutralization test as described previously (Honko *et al*., 2020). Either Omicron (B.1.1.529) or Delta (B.1.617) obtained from BEI Resources (NR-56462 and NR-55673, respectively) was used at 1000 PFU/ml to determine the neutralizing antibody responses in Omicron-infected or Delta-infected groups, respectively. Prior to performing the assay, sera were heat-inactivated at 56°C for 30 minutes and diluted 1 in 50 in DPBS (Gibco, New York, USA), followed by two-fold serial dilutions. The half maximal inhibitory concentration (IC50) for each serum sample was determined from dose response curves constructed in GraphPad Prism 8.4.2.

### Viral Load

Viral loads in hamster nasal turbinates, trachea and lungs were determined by semi-solid Avicel plaque assay as described previously (Honko *et al*., 2020). Briefly, approximately 100 mg of tissue was transferred to a sterile tube (Sarstedt, Nümbrecht, Germany). One milliliter of high glucose Dulbecco’s Modified Eagle Medium (DMEM; Gibco, New York, USA) supplemented with 1 x GlutaMAX-I (Gibco, New York, USA), 1 mM sodium pyruvate (Gibco, New York, USA) and 1 x non-essential amino acids (Gibco, New York, USA) was added to the tube together with a 5 mm stainless steel bead (Qiagen, Maryland, USA). The sample was homogenized for two minutes at 30 Hz using a TissueLyser II (Qiagen, Maryland, USA). The samples were rotated in the adapter and homogenized a second time. Samples were centrifuged at 8 000 x g for two minutes to clarify the supernatant. A 10-fold serial dilution was prepared from each sample from 10^−1^ to 10^−4^. In addition, a 1:1 dilution was prepared from each sample. The dilutions were then plated onto Vero E6 cells seeded at 8 × 10^5^ cells per well (NR-596, BEI Resources). The viral titer in each tissue sample was determined from two replicates of the sample and was reported as PFU/gram tissue. The titer was calculated using the following formula:

Virus titer in PFU/gram = Number of plaques / (virus dilution in well x volume plated in ml)(1000 / weight of tissue in mg)

### Histopathology

The mainstem bronchi of the left lung lobe was clamped with hemostats and removed distally for molecular analysis. Remaining right lung lobes were inflated with sterile DPBS until peak inspiration volume was reached, at which time the distal trachea was transected and both the inflated right lung lobes and nasal passages were immersed in 10% neutral buffered formalin (>10:1 tissue to displaced volume ratio) and inactivated for a minimum of 72 h. Nasal passages were decalcified for 72 hours using Immunocal™ (StatLab, Texas, USA). Inactivated tissues were processed using a Tissue-Tek VIP-5 automated vacuum infiltration processor (Sakura Finetek USA, California, USA) and embedded as formalin fixed paraffin embedded (FFPE) blocks using a HistoCore Arcadia paraffin embedding machine (Leica, Wetzlar, Germany). 5-μm tissue sections were generated using a RM2255 rotary microtome (Leica, Wetzlar, Germany) and were stained with hematoxylin and eosin (H&E) and coverslipped using a ST5010-CV5030 stainer/coverslipper combination (Leica, Wetzlar, Germany). Slides were examined blinded by a board-certified veterinary pathologist (N.A.C.), with subsequent unblinding conducted to generate pathology results. Ordinal scoring criteria and individual animal ordinal scores and qualitative descriptions are described in Supplementary Table 1.

### Statistical Analysis

Data were prepared and analyzed using GraphPad Prism 8.4.2 software (GraphPad Software Inc., California, USA). An unpaired, two-tailed Welch’s t-test was used to determine the significance of differences in the percentage of body weight change, viral load in tissues and neutralizing antibody titers between groups.

## Results

### Clinical observations

The percentage change in animal body weights from the baseline weight recorded prior to inoculation was monitored from the day of challenge up to the day of study termination. Aged animals infected with the Delta variant of SARS-CoV-2 exhibited the greatest percentage weight loss (mean percentage weight change at day 7 of the study = −10%) (Figure 1A, 1C, 1E), followed by young animals infected with the Delta variant (mean percentage weight change at day 7 of the study = −7.8%) (Figure 1A, 1D, 1E). The weight change in both aforementioned groups was statistically significant (p = <0.0001 and p = 0.003 respectively, α = 0.05). In contrast to young animals, aged animals infected with the Delta variant did not completely recover from the weight loss by the end of the study (Figure 1A, 1E). Neither young nor aged animals infected with the Omicron variant of SARS-CoV-2 exhibited a noteworthy change in weight compared to the respective control groups (Figure 1B, 1C, 1D, 1E).

**Figure 1:**
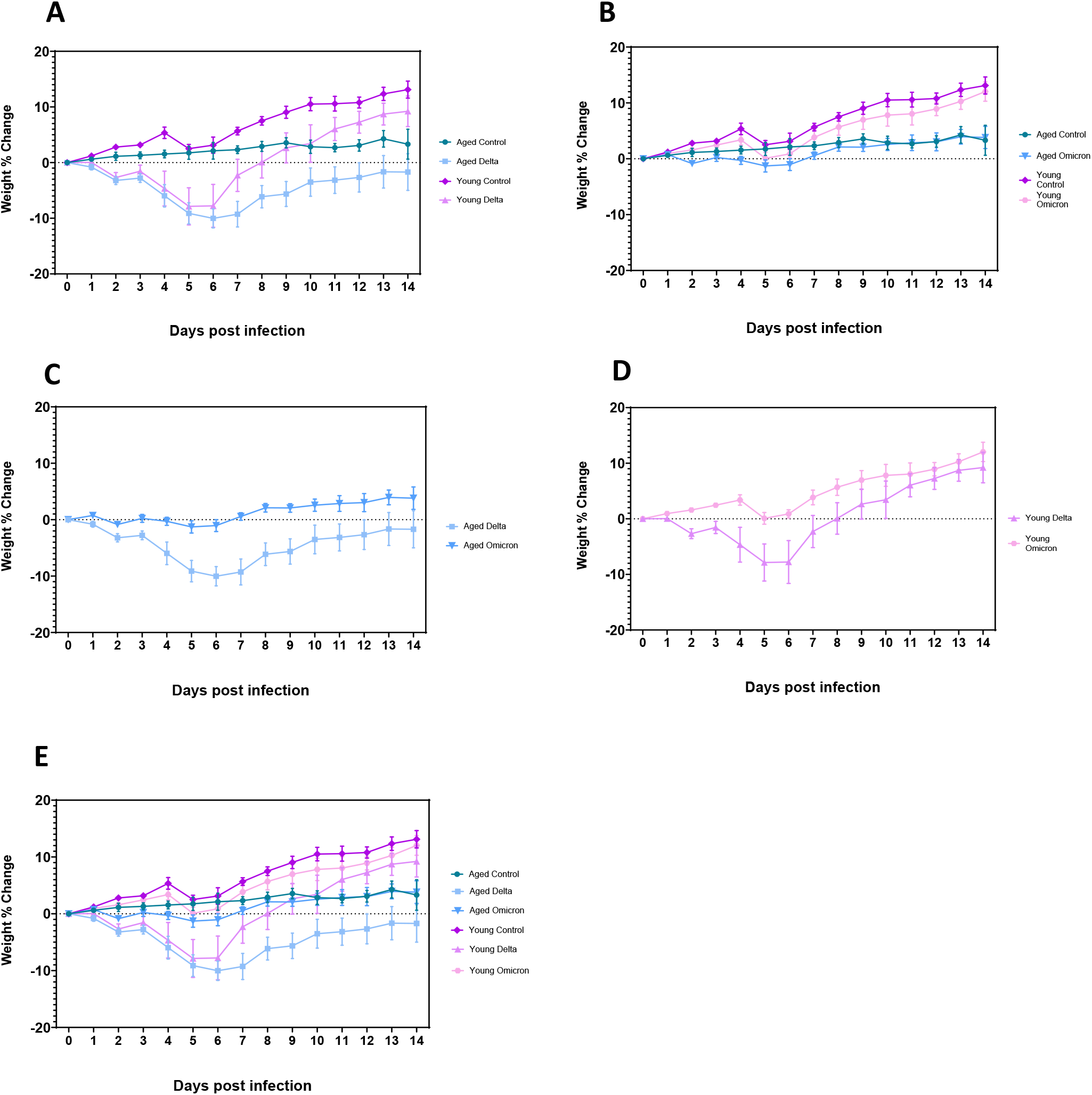
Mean percentage (%) change in animal body weights after challenge with SARS-CoV-2. A) Aged and young hamsters infected with the Delta variant, B) Aged and young hamsters infected with the Omicron variant, C) Aged hamsters infected with either the Delta or Omicron variant, D) Young hamsters infected with either the Delta or Omicron variant, E) Young and aged hamsters infected with either the Delta or Omicron variant.

### Viral load

Viral load in the nasal turbinates, trachea and lungs was determined by plaque assay in a subset of animals in each group euthanized at 3 DPI. Mean SARS-CoV-2 titers in the nasal turbinates of aged and young hamsters infected with the Delta variant were 1.6 × 10^5^ PFU/g and 6.19 × 10^5^ PFU/g, respectively, while the mean SARS-CoV-2 titers in the nasal turbinates of aged and young hamsters infected with the Omicron variant were 2.28 × 10^5^ PFU/g and 3.64 × 10^5^ PFU/g, respectively. There were no statistically significant differences in SARS-CoV-2 titers in the nasal turbinates of any of the infected groups (Figure 2). SARS-CoV-2 titers were lower in the tracheas compared to the lungs in all infected groups, with the mean SARS-CoV-2 titers in the tracheas of aged and young hamsters infected with the Delta variant being 1.32 × 104 PFU/g and 2.09 × 103 PFU/g, respectively, and the mean SARS-CoV-2 titers in the tracheas of aged and young hamsters infected with the Omicron variant being 5.76 × 103 PFU/g and 1.38 × 103 PFU/g, respectively. There was a statistically significant difference in SARS-CoV-2 titers in the tracheas of aged and young hamsters infected with the Delta variant (p = 0.04, α = 0.05) with mean tracheal SARS-CoV-2 titers being higher in the aged group (Figure 3). However, there was no significant difference in tracheal SARS-CoV-2 titers in young and aged hamsters infected with the Omicron variant, or in the tracheas of young or aged hamsters infected with either the Delta or the Omicron variant.

**Figure 2:**
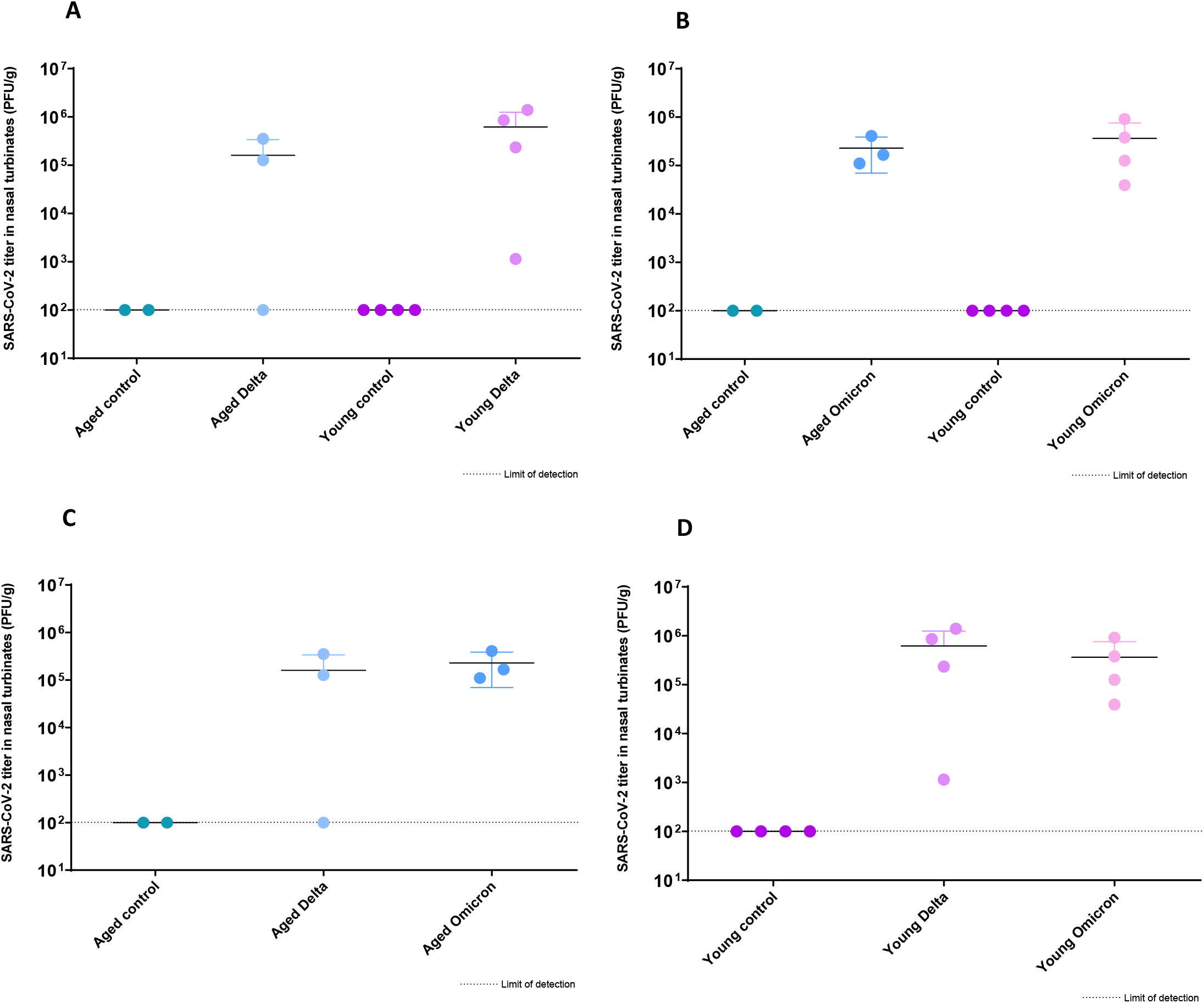
SARS-CoV-2 titer in the nasal turbinates of hamsters challenged with one of two SARS-CoV-2 variants of concern. A) Delta-infected hamsters, B) Omicron-infected hamsters, C) Aged hamsters, D) Young hamsters. The graphs represent the mean SARS-CoV-2 titer in the tissue from each group, with error bars representing the standard deviation.

**Figure 3:**
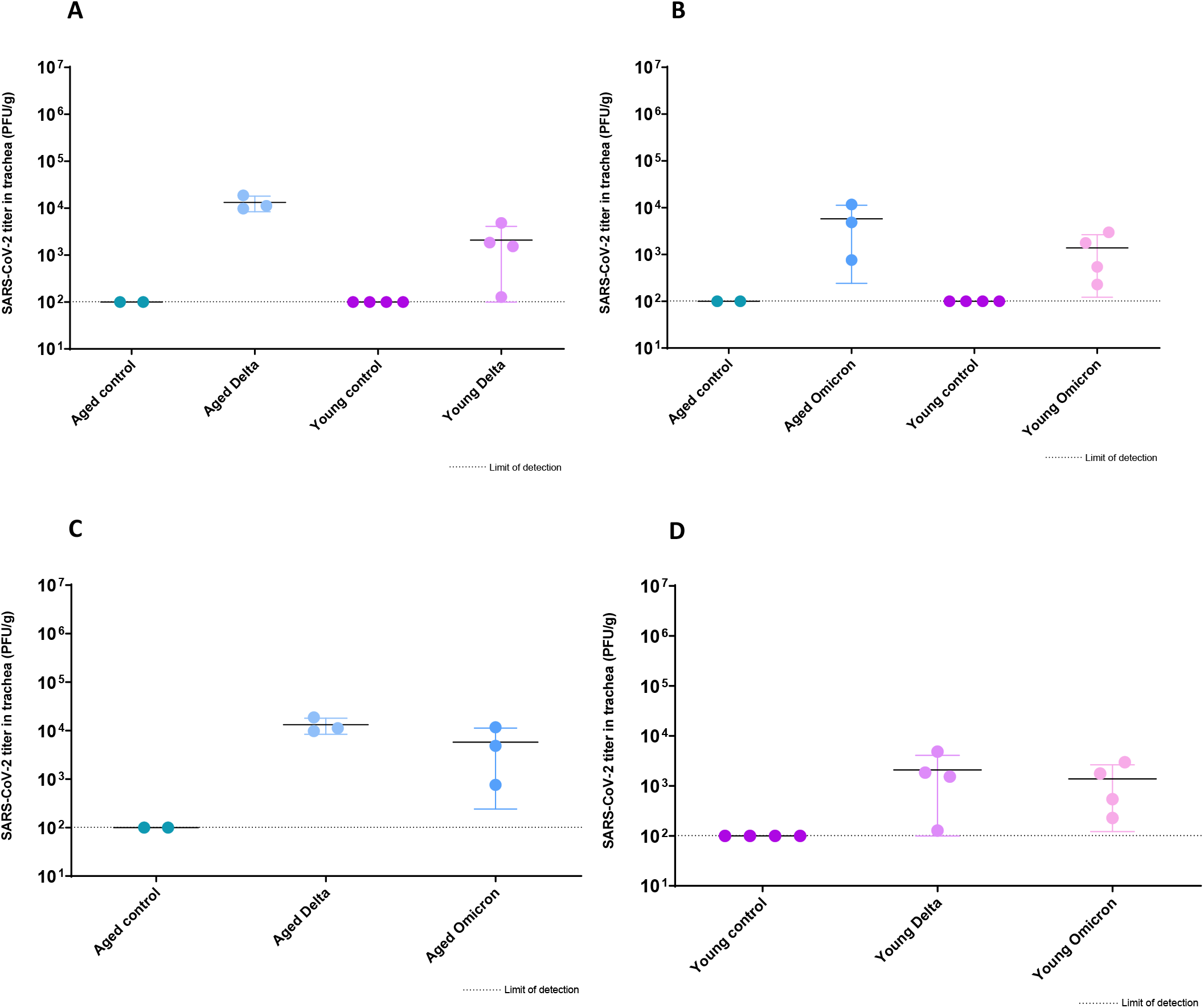
SARS-CoV-2 titer in the tracheas of hamsters challenged with one of two SARS-CoV-2 variants of concern. A) Delta-infected hamsters, B) Omicron-infected hamsters, C) Aged hamsters, D) Young hamsters. The graphs represent the mean SARS-CoV-2 titer in the tissue from each group, with error bars representing the standard deviation.

The mean SARS-CoV-2 titers in the lungs of aged and young hamsters infected with the Delta variant were 1.76 × 10^6^ PFU/g and 2.91 × 10^6^ PFU/g, respectively, while the mean SARS-CoV-2 titers in the lungs of aged and young hamsters infected with the Omicron variant were 4.21 × 10^5^ PFU/g and 1.43 × 10^5^ PFU/g, respectively. There was no significant difference in SARS-CoV-2 titers in the lungs of aged and young hamsters infected with the Delta variant, or in the lungs of aged and young hamsters infected with the Omicron variant. There was a statistically significant difference in the SARS-CoV-2 titers in the lungs of young hamsters infected with either the Delta or the Omicron variant (p = 0.006, α = 0.05) with higher mean SARS-CoV-2 lung titers in the young Delta infected group, but not in aged hamsters infected with either of the two variants (Figure 4).

**Figure 4:**
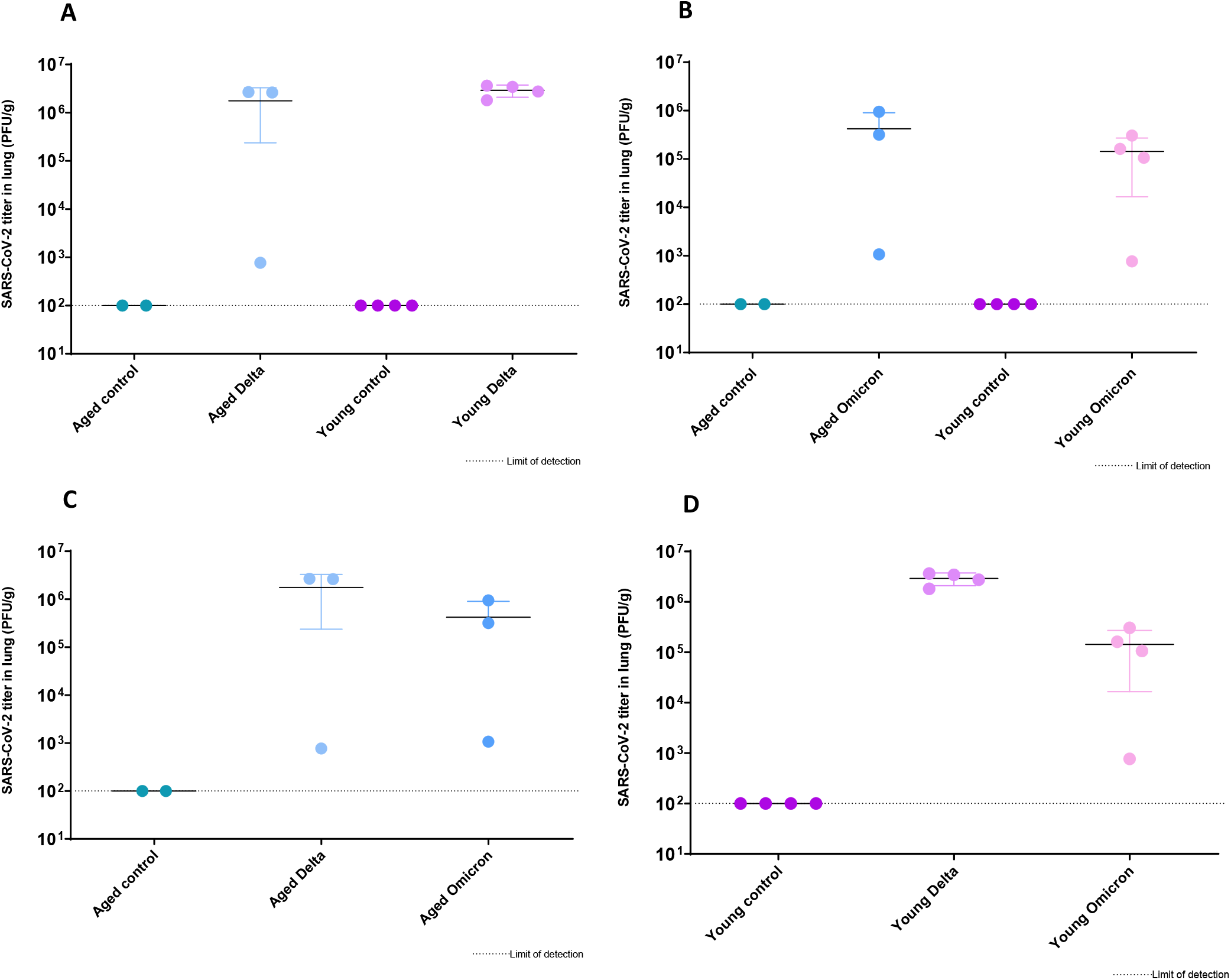
SARS-CoV-2 titer in the lungs of hamsters challenged with one of two SARS-CoV-2 variants of concern. A) Delta-infected hamsters, B) Omicron-infected hamsters, C) Aged hamsters, D) Young hamsters. The graphs represent the mean SARS-CoV-2 titer in the tissue from each group, with error bars representing the standard deviation.

### Neutralizing antibody titers

The mean half maximal inhibitory concentration (IC50) of neutralizing antibodies to SARS-CoV-2 was determined for each group from sera collected on day 14 PI. Figure 5 shows the IC50 for each group of animals challenged with either DPBS, SARS-CoV-2 Delta variant or SARS-CoV-2 Omicron variant, with error bars representing the standard deviation from the mean. There was a significant difference in neutralizing antibody titers produced in young versus aged animals infected with the Omicron variant (p=0.05, α = 0.05), with young animals producing higher mean IC50s (mean aged IC50 = 1384.1, mean young IC50 = 3831.3). Although the mean IC50s were also higher in young animals infected with the Delta variant compared to the aged group, the difference was not statistically significant. Mean IC50s produced by infected animals were higher against the Delta variant than the Omicron variant, with a significant difference in titers produced against the two variants in the aged group (p = 0.004, α = 0.05).

**Figure 5:**
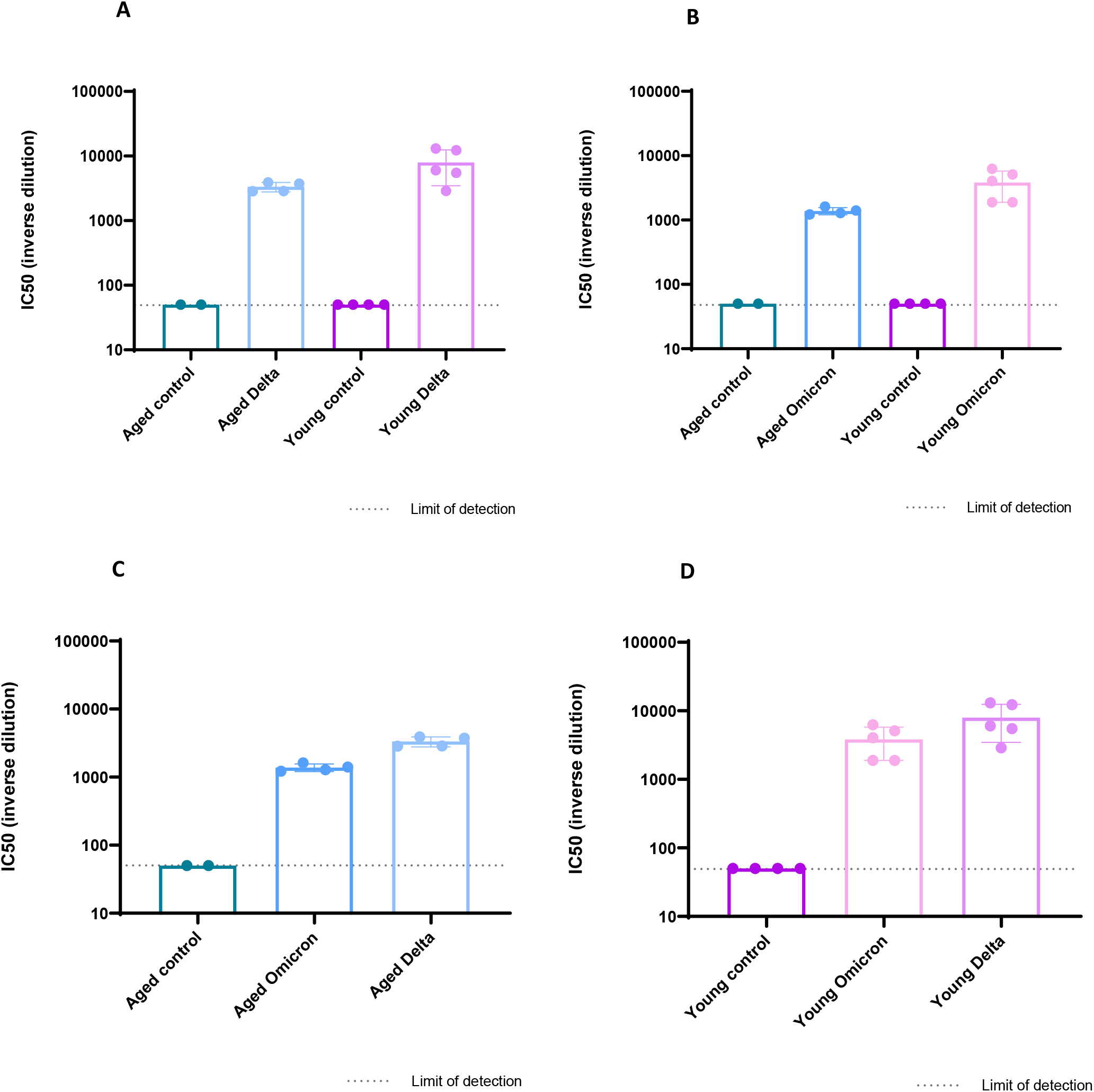
Mean half maximal inhibitory concentration of neutralizing antibodies in sera collected from animals 14 days post inoculation with one of two SARS-CoV-2 variants of concern. A) Delta-infected hamsters, B) Omicron-infected hamsters, C) Aged hamsters, D) Young hamsters. Error bars represent the standard deviation from the mean.

### Histopathology

#### Irrespective of age, Syrian hamsters inoculated with the Delta variant results in severe necrotizing pan-rhinitis, while the Omicron variant acutely causes rhinitis with epithelial injury and noteworthy olfactory epithelium (OE) sparing

Post-fixation, whole heads were decalcified and processed as a midline sagittal section affording simultaneous examination of the respiratory epithelium (RE), olfactory epithelium (OE) and olfactory bulb (OB). Nasal passage ordinal scores and representative images of Sham/DPBS, Delta, and Omicron inoculated hamsters at 3 DPI are represented in Figures 6 and 7. At 3 DPI, both Delta and Omicron inoculated hamsters, irrespective of age, displayed similar histopathologic patterns in the RE compartment of the nasal cavity. This was characterized by moderate to regionally severe neutrophilic inflammation and edema of the lamina propria, with moderate to severe degeneration and necrosis of the RE, characterized by loss of cilia, cell rounding with hypereosinophilic cytoplasm, and pyknotic nuclei. Additionally, RE segmentally displayed squamous metaplasia. Glands and ducts within the RE lamina propria were multifocally necrotic or dilated and lined by attenuated epithelium with accumulated luminal neutrophilic infiltrate respectively.

**Figure 6:**
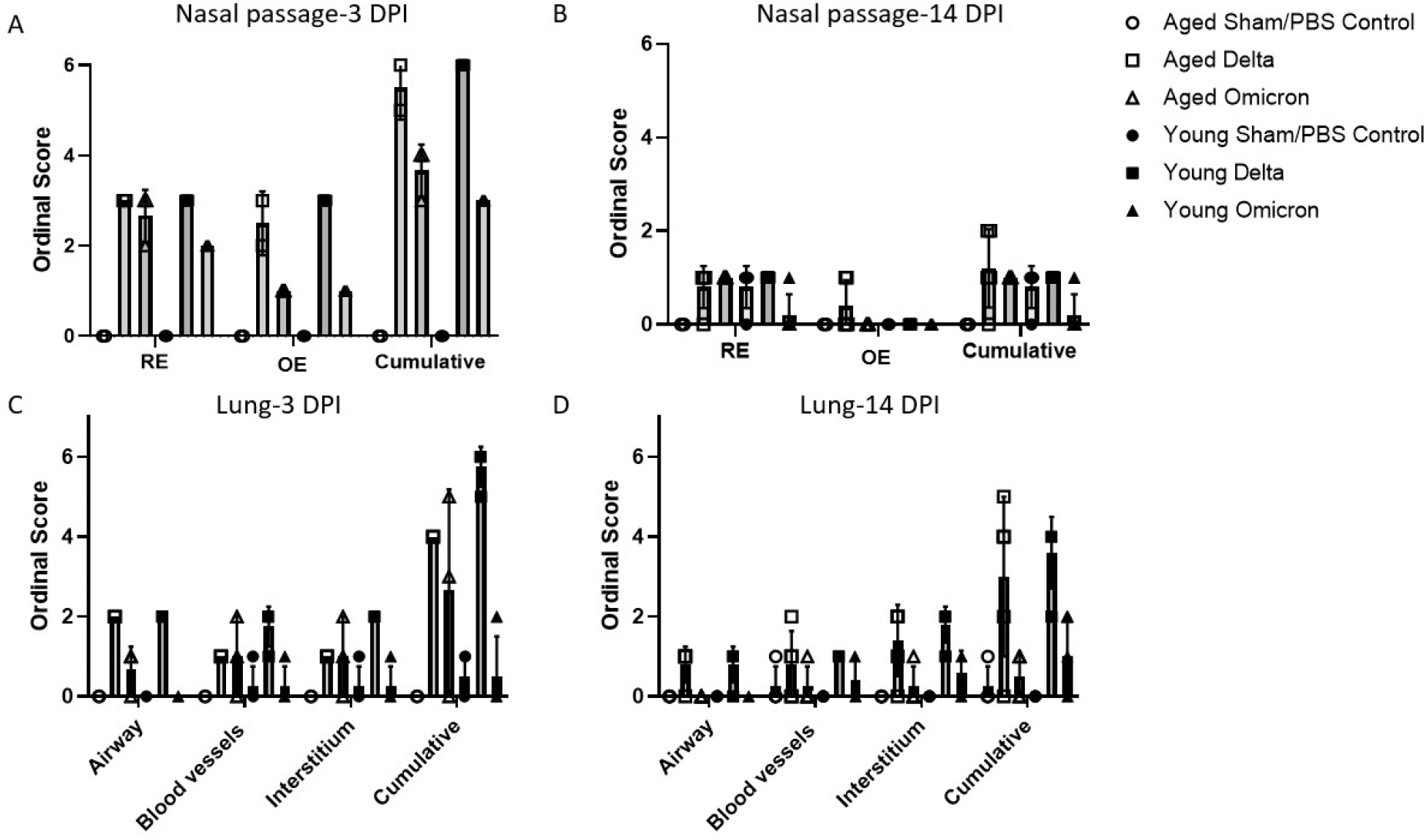
Mean nasal passage and lung ordinal scores from animals 3- and 14-days post inoculation (DPI) with one of two SARS-CoV-2 variants of concern. A) Nasal passage, 3 DPI. B) Nasal passage, 14 DPI. C) Lung, 3 DPI. D) Lung, 14 DPI. Error bars represent the standard deviation from the mean.

**Figure 7:**
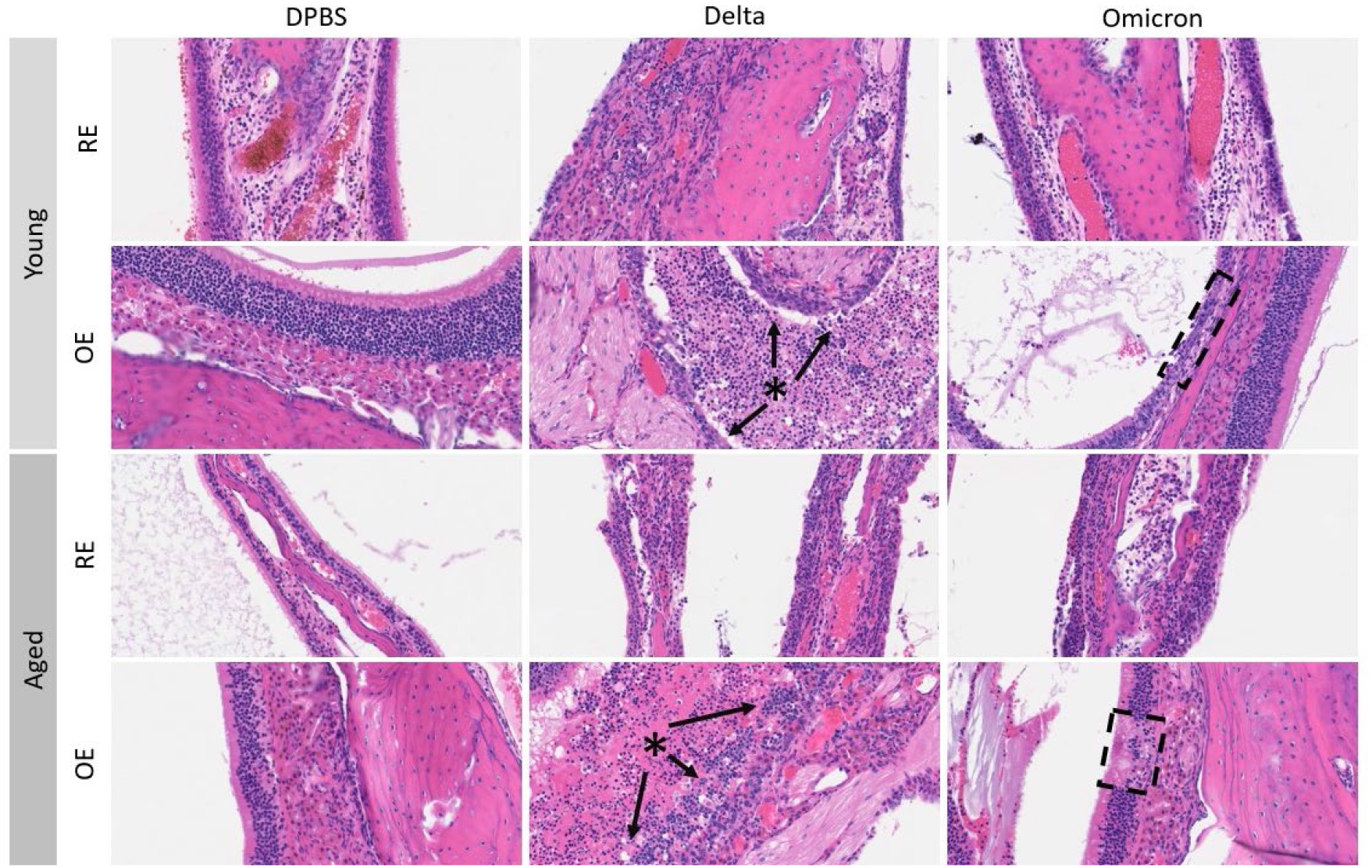
Representative hematoxylin and eosin (H&E) images of the respiratory epithelium (RE) and olfactory epithelium (OE) at day 3 post-inoculation (DPI) in young and aged Syrian hamsters infected with Sham/DPBS or the Delta or Omicron variant of SARS-CoV-2. Loss of OE pseudostratification (arrows) and accumulation of necrotic debris and inflammatory infiltrate in neighboring nasal meatuses (asterisks) was severe in all animals receiving Delta variant, while milder and more localized OE injury was observed with Omicron (hashed boxes). 200x (OE) or 400x (RE) total magnification.

Distinctly, hamsters inoculated with Delta variant, again irrespective of age, exhibited severe, and at times near global necrosis extending to the level of basal cells, with otherwise complete dissolution of the normal pseudostratified columnar layering of olfactory neurons and sustentacular cells. Furthermore, the neighboring nasal meatuses overlying the OE were flooded by severe neutrophilic infiltrate, denuded OE, necrotic debris, and proteinaceous exudate. In contrast, mild segmental OE degeneration and necrosis evidenced by loss of pseudostratification with absence of exudate in neighboring nasal meatuses was observed in Omicron inoculated hamsters, with absent to mild neutrophilic infiltrate. Nasal passages of DPBS control groups were within normal limits; furthermore, the olfactory bulb (OB) was histologically normal in all hamsters irrespective of cohort, histomorphologically suggesting a lack of neuroinvasion.

#### Irrespective of age, near complete restoration of the nasal passage occurs by 14 DPI in both Delta and Omicron inoculated Syrian hamsters

Nasal passage ordinal scores and representative images of Sham/DPBS, Delta, and Omicron inoculated hamsters at 14 DPI are represented in Figures 6 and 8. At 14 DPI, residual degeneration and necrosis of RE and OE were no longer evident, although mild regional neutrophilic infiltrate within the lamina propria of the RE and less commonly OE were observed; however, mild neutrophilic infiltrate was also observed in some of the Sham/DPBS-inoculated hamsters suggesting this observation could in part reflect an incidental background finding. The restoration of nasal mucosal surfaces in the RE and OE supports a robust reparative response. Notably, one hamster in the aged Delta group displayed focally extensive OE dysplasia adjacent to the cribriform plate, as evidenced by localized formation of rosettes admixed with variable amounts of residual glands, nerve fascicles, and adipocytes that locally replaced the nasal meatus (Supplemental Figure 1). This finding suggests that albeit not common, long-term aberrations in the OE are feasible in Syrian hamsters that could result in long term deficiencies to olfaction.

**Figure 8:**
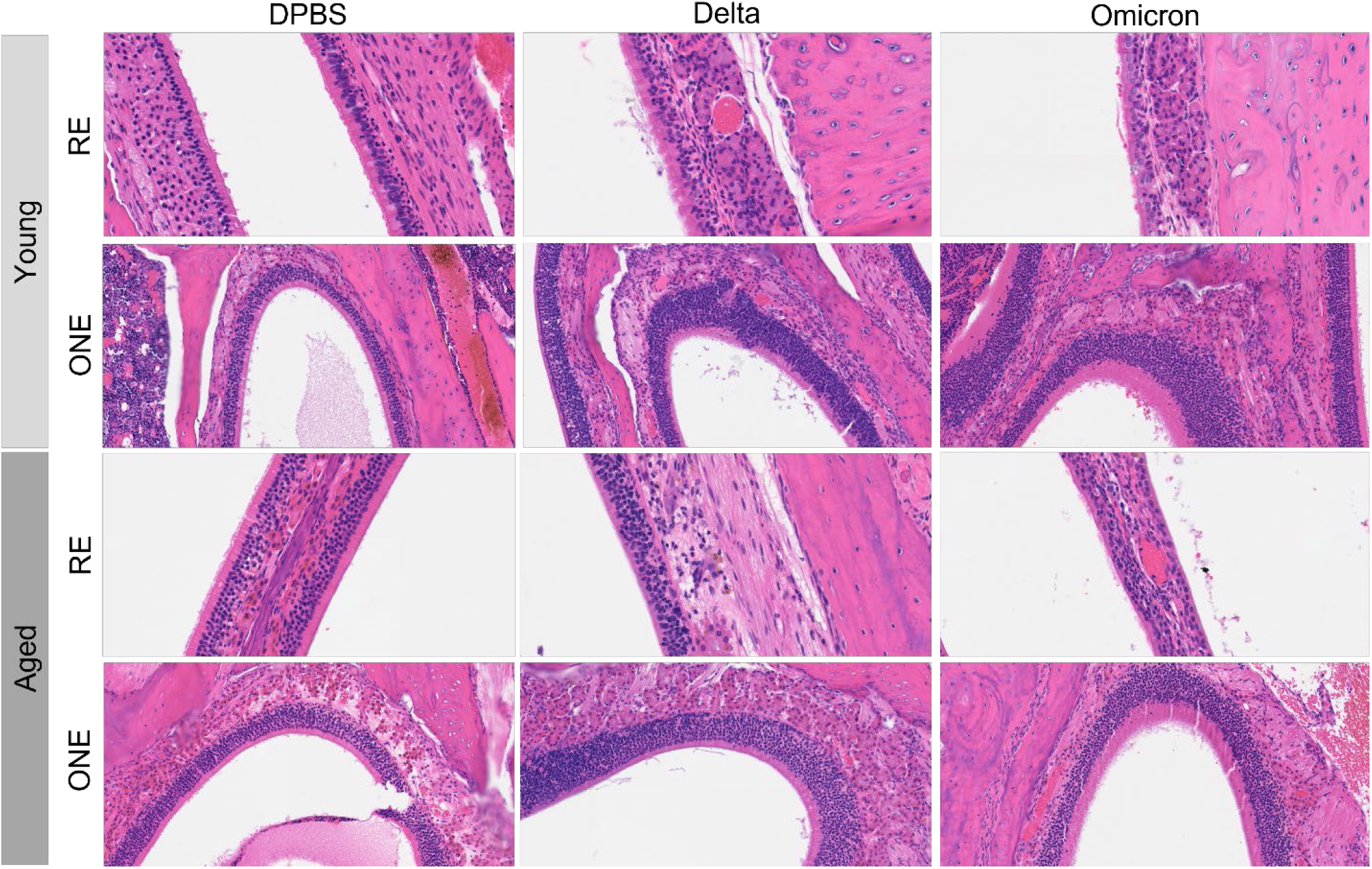
Representative hematoxylin and eosin (H&E) images of the respiratory epithelium (RE) and olfactory epithelium (OE) at day 14 post-inoculation (DPI) in young and aged Syrian hamsters infected with Sham/DPBS, Delta, or Omicron variant of SARS-CoV-2. 200x (OE) or 400x (RE) total magnification.

#### Irrespective of age, Omicron variant acutely causes mild pulmonary disease, while Delta variant results in moderate acute necrotizing bronchointerstitial pneumonia

Lung ordinal scores and representative images of Sham/DPBS, Delta, and Omicron at 3 DPI are represented in Figures 6 and 9. At 3 DPI most Omicron-inoculated aged and young hamsters histologically lacked or displayed mild localized bronchiole degeneration, necrosis, and peribronchiolar inflammatory infiltrate, which was invariably present multifocally and of moderate severity in Delta-inoculated hamsters irrespective of age. Similarly, with rare exception, pulmonary parenchyma of aged and young Omicron inoculated animals was either histologically normal or displayed mild mononuclear perivascular and/or interstitial infiltrate with absence of alveolar hemorrhage, fibrin, or edema. In comparison, pulmonary parenchyma including alveolar septa, alveoli, and perivascular compartments in Delta-inoculated aged and young hamsters routinely contained mild to moderate multifocal mononuclear +/-neutrophilic infiltrate, with sporadic alveolar hemorrhage, edema, and fibrin polymerization.

**Figure 9:**
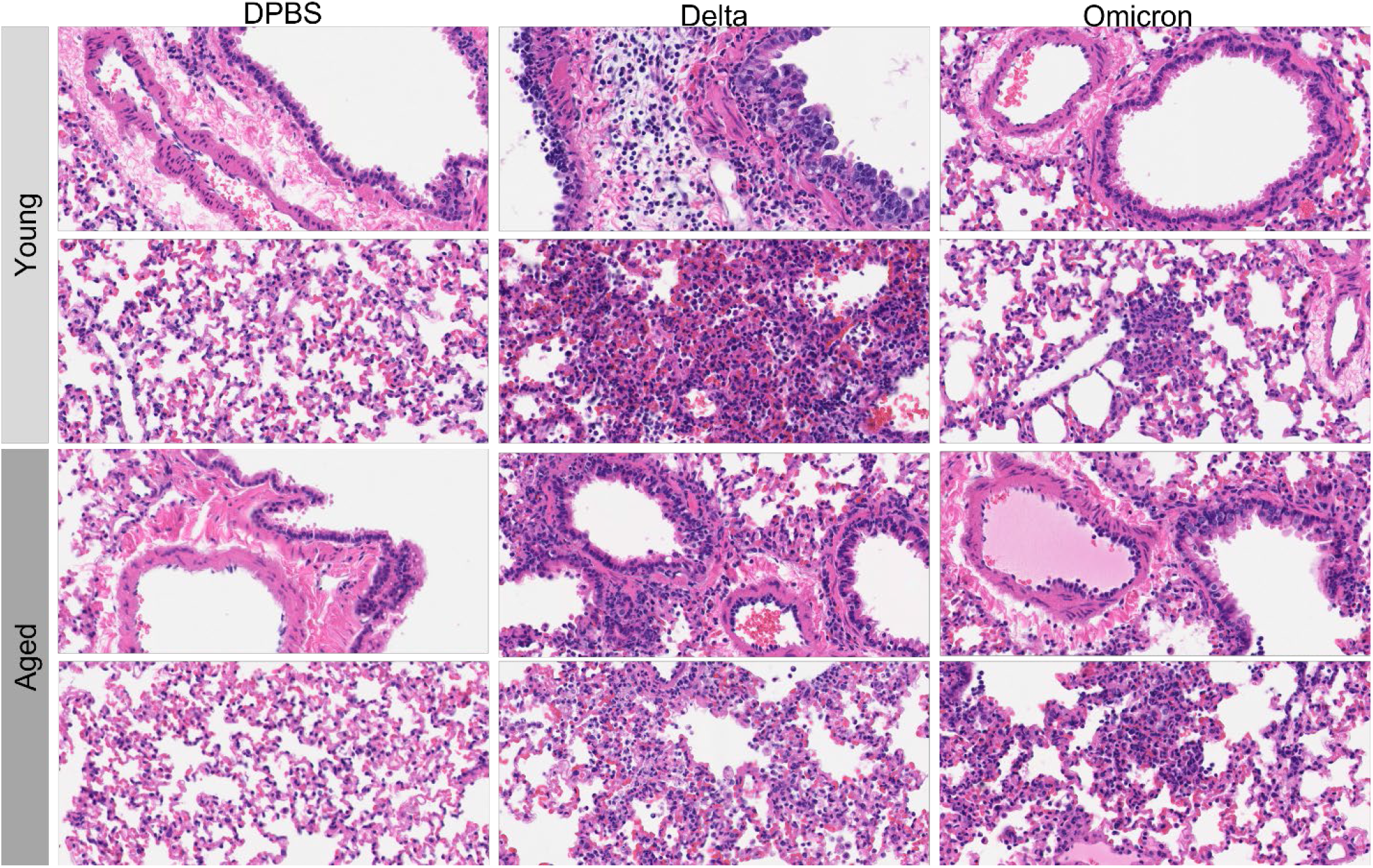
Representative hematoxylin and eosin (H&E) images of the lung at 3 days post-inoculation (DPI) in young and aged Syrian hamsters infected with Sham/DPBS, Delta, or Omicron variant of SARS-CoV-2. Bronchiolar degeneration and necrosis, as well as evidence of alveolar damage (i.e., fibrin polymerization, hemorrhage, edema and neutrophil influx) were more frequent and severe in Delta infected hamsters regardless of age. 400x total magnification.

#### Irrespective of age, by 14 DPI residual pulmonary pathology was absent to mild in Omicron-inoculated Syrian hamsters, with increased evidence of epithelial repair in Delta-inoculated hamsters consistent with more severe prior lung injury

Lung ordinal scores and representative images of Sham/DPBS, Delta, and Omicron inoculated hamsters at 14 DPI are represented in Figures 6 and 10. Mild segmental bronchiolar hyperplasia and mild peribronchiolar mononuclear infiltrate were observed exclusively in hamsters inoculated with Delta, reflective of prior necrotizing bronchiolitis that was observed acutely (3 DPI). Furthermore, alveolar type 2 (AT2) hyperplasia and alveolar bronchiolization were either not observed or mild and focal in Omicron-inoculated hamsters, while these findings were routinely observed in Delta-inoculated hamsters. Additionally, mild to moderate lymphohistiocytic perivascular and interstitial infiltrate and alveolar histiocytosis with sporadic pigment (presumed hemosiderin) were more frequently observed in Delta-inoculated hamsters consistent with prior alveolar damage and hemorrhage. Plural fibrosis was sporadically observed in both Omicron and Delta cohorts, irrespective of age, but was at worst only mild and segmental. Taken together, histopathologic findings are consistent with more severe prior acute lung injury induced by Delta variant when compared to Omicron and a more robust affiliated reparative response.

**Figure 10:**
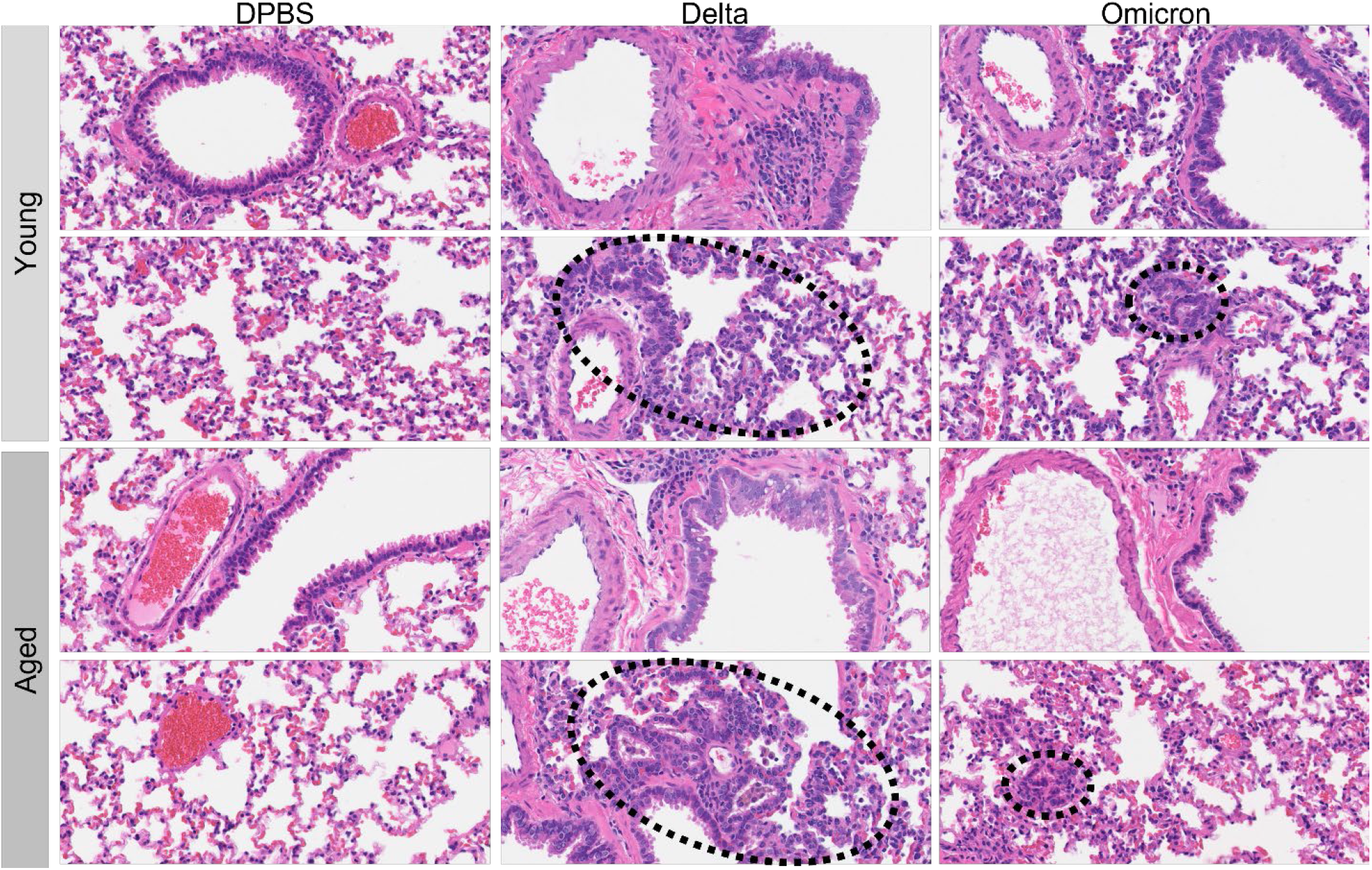
Representative hematoxylin and eosin (H&E) images of the lung at 14 days post-inoculation (DPI) in young and aged Syrian hamsters infected with Sham/DPBS, Delta, or Omicron variant of SARS-CoV-2. Alveolar type 2 (AT2) hyperplasia and/or bronchiolization of alveoli (reparative responses) were readily observed in Delta infected hamsters compared to Omicron regardless of age suggestive of more severe prior lung injury (hashed ovals). 400x total magnification.

## Discussion

Although the SARS-CoV-2 Omicron variant has by far caused the most confirmed cases of COVID-19 to date, this variant is markedly attenuated, with a 73% reduced risk of hospitalization having been reported when compared to Delta (Veneti *et al*., 2022). Furthermore, some common COVID-19 signs such as anosmia and ageusia have been reported at markedly lower prevalence (i.e., 8.3% and 9%) compared to pre-Omicron reports (38.2% & 36.6%) (Mutiawati *et al*., 2021; Maisa *et al*., 2022). Syrian hamsters are a well-established animal model that have proven to be an excellent platform to evaluate efficacy of vaccines and therapeutics for SARS-CoV-2 and mimic many clinicopathologic hallmarks of moderate self-limiting disease in humans (Imai *et al*., 2020; Gruber *et al*., 2021). However, this model is not without limitations. It has been established that severe COVID-19 disease is associated with one or more co-morbidities (Zhou *et al*., 2020) which is not readily reproducible in animals, representing a challenge in developing models that fully represent the more severe manifestation of the disease.

Our findings illustrate that weight loss was more pronounced and recovery incomplete in aged animals infected with the Delta variant compared to young animals. These findings are consistent with a previous study comparing infection with an early variant of SARS-CoV-2 (BetaCoV/Germany/BavPat1/2020) in aged and young Syrian hamsters (Osterrieder *et al*., 2020) where weight loss was more distinct in older animals. Infection with Omicron did not result in weight loss in animals in either cohort. Omicron replicated to similar levels as Delta in the upper respiratory tract (nasal turbinates and trachea) with a trend towards decreased viral load in the lower respiratory tract (lungs) during acute disease (3 DPI), with no significant differences in tissue viral loads of aged versus young animals for either variant. In contrast, histopathologic differences were clear among the Omicron and Delta variants in both the nasal passages and lungs. Specifically, severe necrosis was observed acutely (3 DPI) in the OE of Delta-infected animals, while this phenotype was either absent or mild and segmental in Omicron-infected animals regardless of age. Acute Omicron infection (3 DPI) also resulted in either no evident pathology or mild pulmonary disease in both young and aged animals compared to uniformly moderate acute necrotizing bronchointerstitial pneumonia observed in Delta-infected animals at 3 DPI.

At 14 DPI, with the exception of localized OE dysplasia in one aged hamster attributed to dysfunctional repair of prior severe injury, there was histologic resolution of disease in the nasal passages of all animals consistent with previous reports in Syrian hamsters infected with BetaCoV/Hong Kong that causes severe OE necrosis acutely (Sia *et al*., 2020). These findings are consistent with the transient albeit variable loss of smell reported in humans with an increased prevalence in Delta compared to Omicron; however, these also suggest a mechanism by which long-term aberrations in the OE could result in long term olfaction dysfunction with variants that cause severe OE necrosis.

Similarly in the lung at 14 DPI, much of the acute pathology observed at 3 DPI (necrotizing bronchiolitis and alveolar injury in Delta animals) had resolved, with residual findings attributable to reparative responses stemming from prior acute lung disease, all of which were either absent or rarely observed in animals inoculated with the Omicron variant.

Advanced age is reported as a risk factor for development of severe and lethal COVID-19; however, this has not been consistently reproduced in Syrian hamsters. Reports in support of this notion documented a reduced capacity to regain body weight and delayed viral clearance (Osterrieder *et al*., 2020; Griffin *et al*., 2021). While we also observed the former with Delta, we cannot comment on the latter as samples for viral assays were collected at 3 DPI and 14 DPI, which represents temporal peak and resolution of detectable virus in Syrian hamsters. Another report refutes age as a key determinant of disease outcome, documenting an absence of outcome on severity or course of infection (Rosenke *et al*., 2020).

Our results clearly indicate an attenuated clinicopathologic outlook of SARS-CoV-2 related disease in Syrian hamsters infected with the Omicron variant compared to the Delta variant irrespective of age that is much less apparent based on tissue viral loads. Taken together, these findings are consistent with the reported increased transmissibility of Omicron variant, with reduced risk for hospitalization. Acknowledging comparable viral loads with decreased pathogenicity, its logical to consider that sub-clinically infected humans are more likely to unknowingly infect others by not participating in preventative public health measures such as isolation. These findings may assist in informing vaccine and therapeutic development strategies for SARS-CoV-2 variants of concern.

## Acknowledgements

The authors would like to thank Madison Pacaro, Igor Gavrish, Aoife O’Connell, Hans P. Gertje and the Animal Research Services team of the National Emerging Infectious Diseases Laboratories of Boston University for technical assistance with this experiment. Histopathology figures were generated using a whole slide scanner acquired from a NIH SIG award (S10OD030269-N.A.C.).

## Author Contributions

Nadia Storm, Nicholas A. Crossland and Anthony Griffiths designed the experiment. Nadia Storm, Nicholas A. Crossland and Lindsay McKay performed the experiment. Nadia Storm, Nicholas A. Crossland and Anthony Griffiths analyzed the data. Nadia Storm and Nicholas A. Crossland wrote the paper. Anthony Griffiths supervised and acquired funding for the experiment. All authors revised the draft paper. This research was supported by Massachusetts Consortium on Pathogen Readiness.

---

**Supplemental Figure 1:**
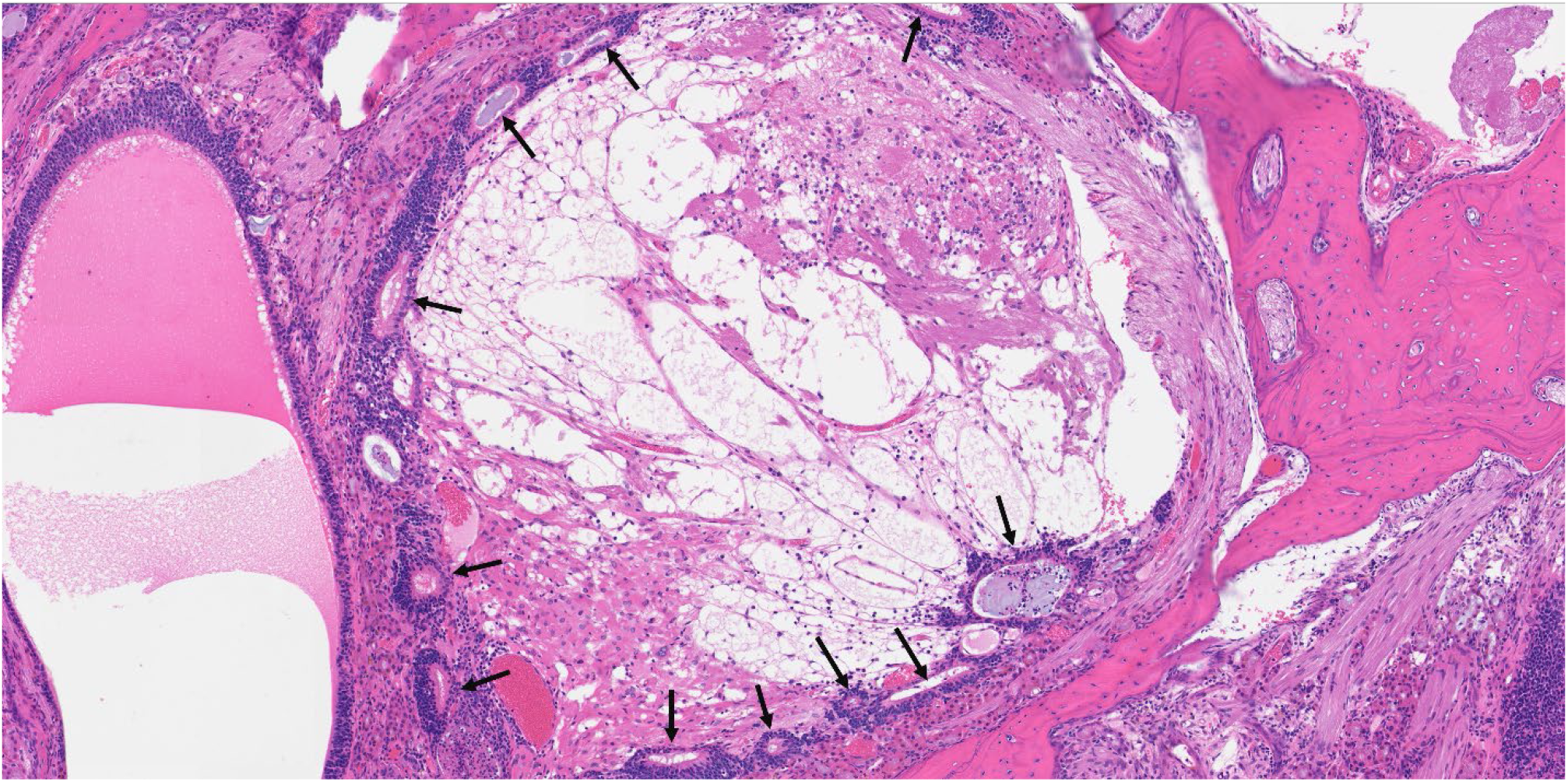
Hematoxylin and eosin (H&E) image of aged Delta inoculated Syrian hamster, 14 days post-inoculation (DPI). Focally extensive olfactory epithelium (OE) dysplasia adjacent to the cribriform plate. Peripheralized OE rosettes (arrows) occasionally dilated by necrotic debris and neutrophils, with replacement of the nasal meatus by haphazardly arranged aggregates of glands, adipocytes, and nerve fascicles. 100x total magnification.

**Table S1.**
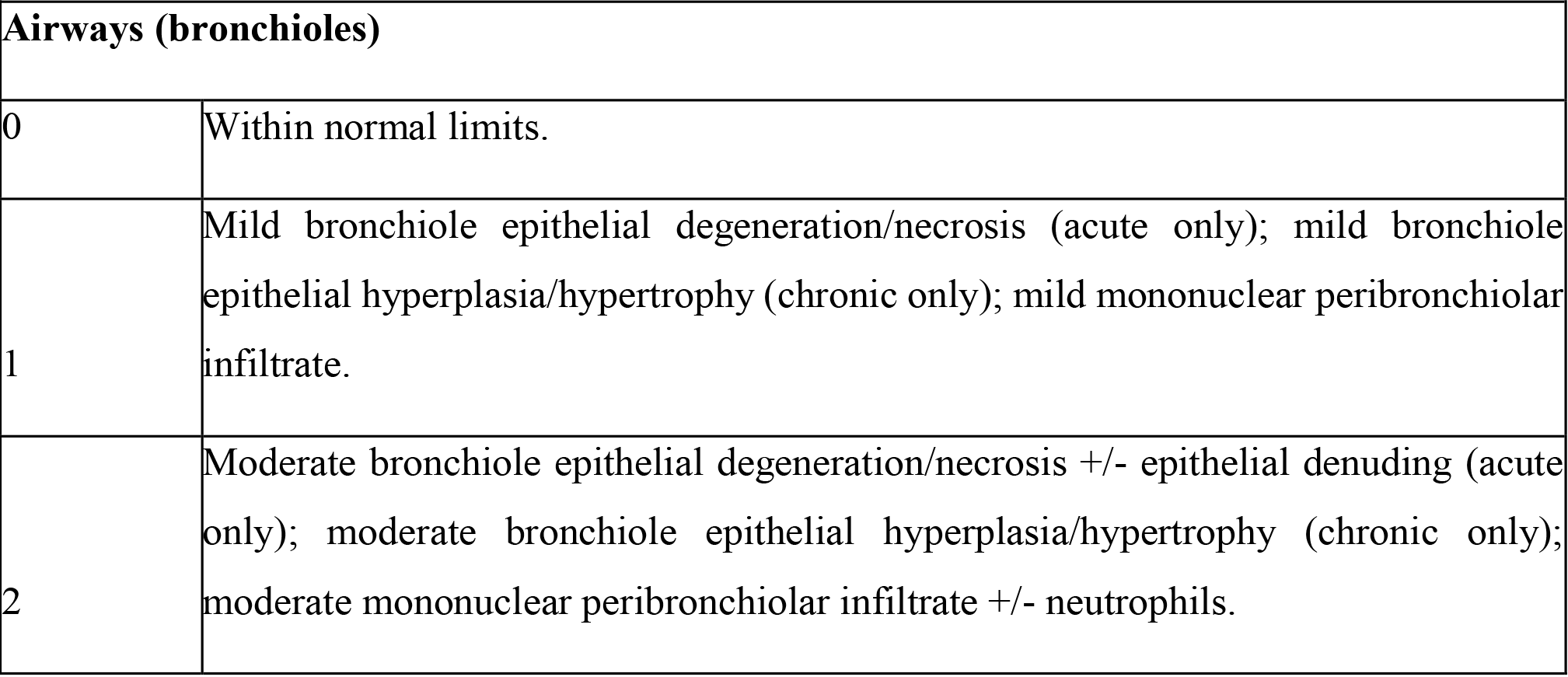
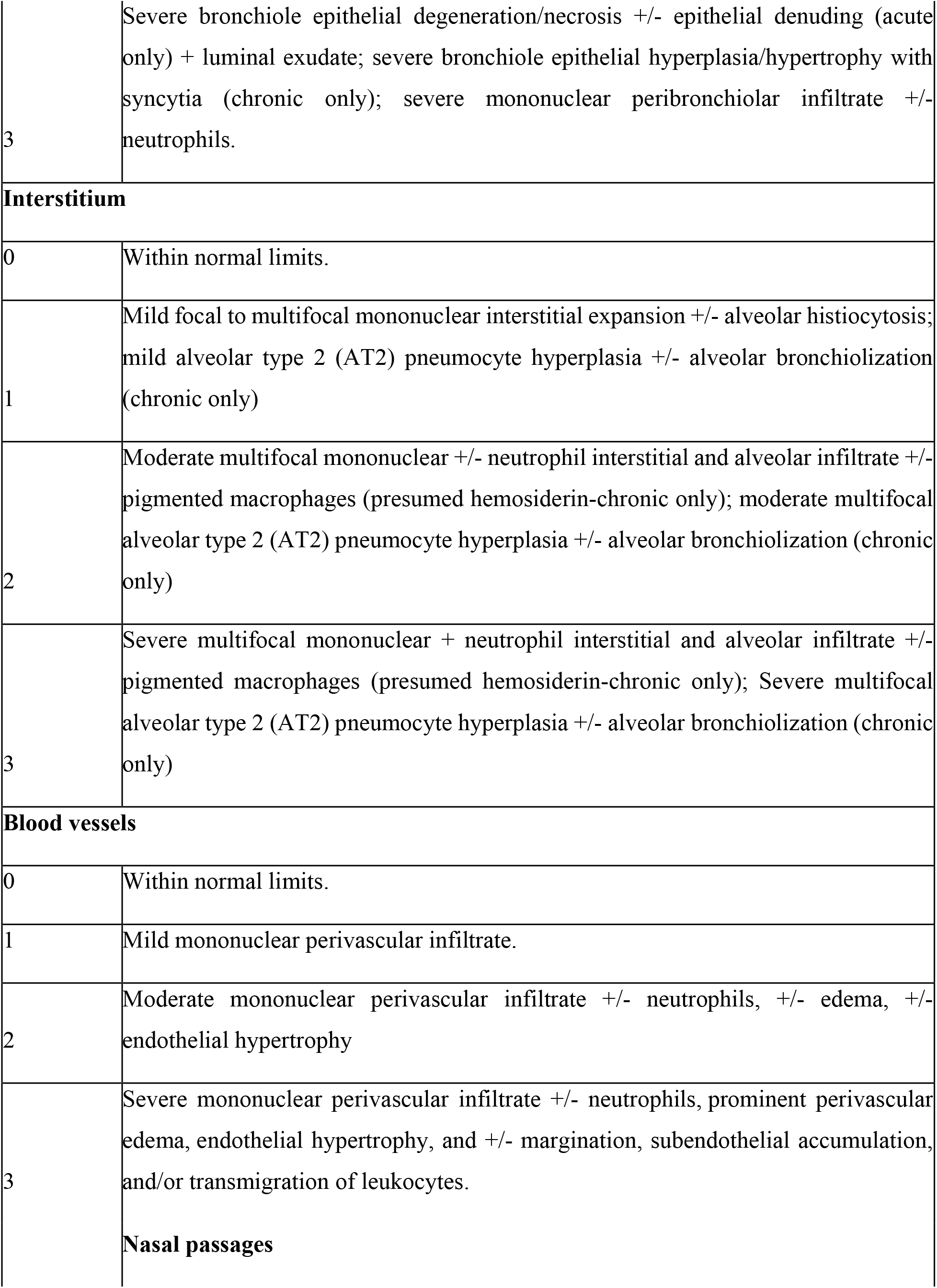
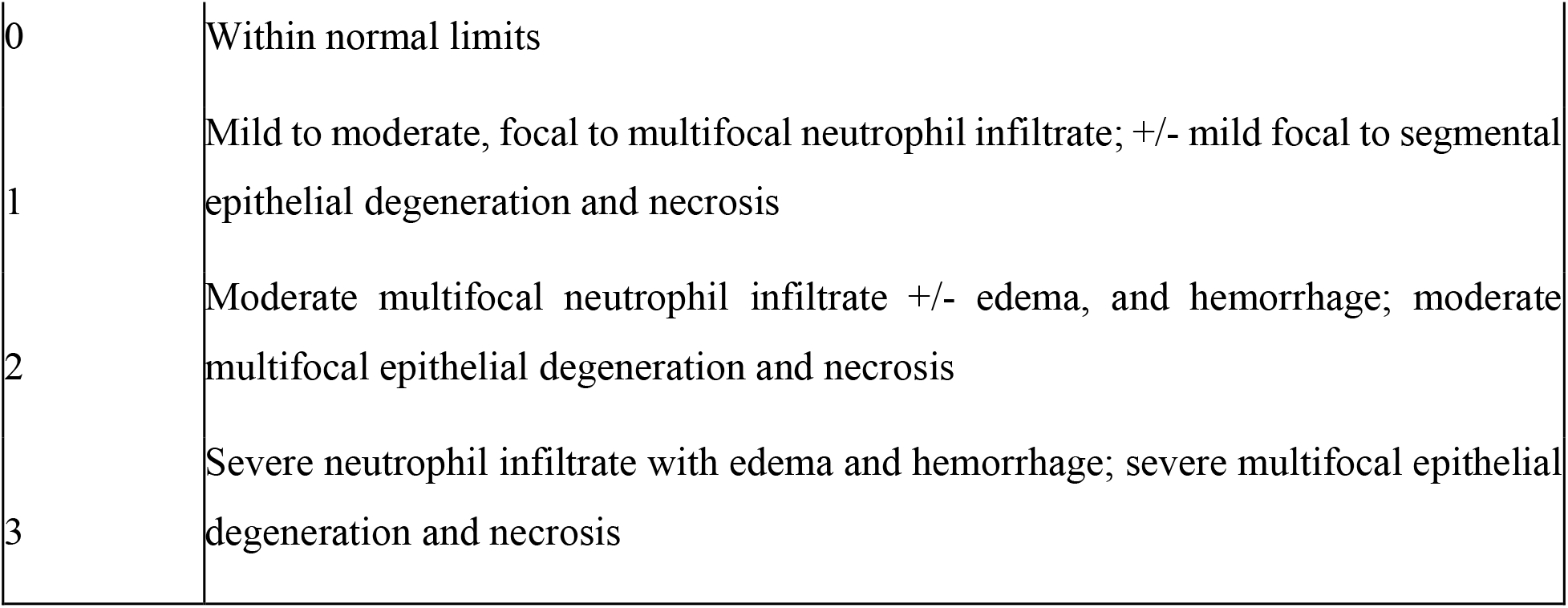
Ordinal Histopathology Criteria.

